# Bi-directional communication between monocytes and trophoblasts under hypoxia and hypoxia-reperfusion conditions

**DOI:** 10.1101/2023.09.23.558721

**Authors:** Hannah Yankello, Yerim Lee, Christina Megli, Elizabeth Wayne

## Abstract

**Introduction:** Pregnancy-related disorders such as preeclampsia are associated with syncytiotrophoblast (STB) stress and monocyte dysregulation. It remains unclear whether this stress derives from prolonged placental hypoxia or a hypoxia-reperfusion-type injury. Thus, this study investigated how these two models of STB stress impact trophoblast-monocyte interactions.

**Method:** Cobalt chloride chemically induced hypoxia in BeWo b30 cells. A transwell coculture system was used to examine trophoblast-monocyte signaling. qPCR quantified gene expression changes following coculture. Monocyte phagocytosis of *E. Coli* or adhesion to placental cells was determined via flow cytometry. Monocyte migration to placental signals was quantified using a cell counter.

**Results:** Cobalt chloride induced a hypoxic state in BeWo b30s. Reperfusion restored the expression of indirect hypoxia genes and ER stress genes. Coculturing THP-1 monocytes with normoxic, hypoxic, and hypoxic-reperfused BeWo b30s promoted b30 survival but not wound-healing capacity. Compared to hypoxic-reperfused BeWos, hypoxic cells increased monocyte adhesion and inflammatory gene expression, decreased monocyte phagocytosis, and did not change monocyte migration. Finally, placental signaling in early-onset PE decreased monocyte chemotaxis, but monocyte precondition more strongly influenced migration compared to placental state.

**Discussion:** Overall, hypoxic placental signals most effectively recapitulate monocyte functional behavior observed in preeclampsia. Further research is needed to understand spatial and temporal changes in monocyte-trophoblast interactions and pregnancy outcomes. Monocyte chemotaxis to primary placental signals varied by gestational age, maternal diagnosis, and monocyte condition, implying monocytes could be used as functional biomarkers to predict their behavior at the maternal-fetal interface as well as the onset of disease.

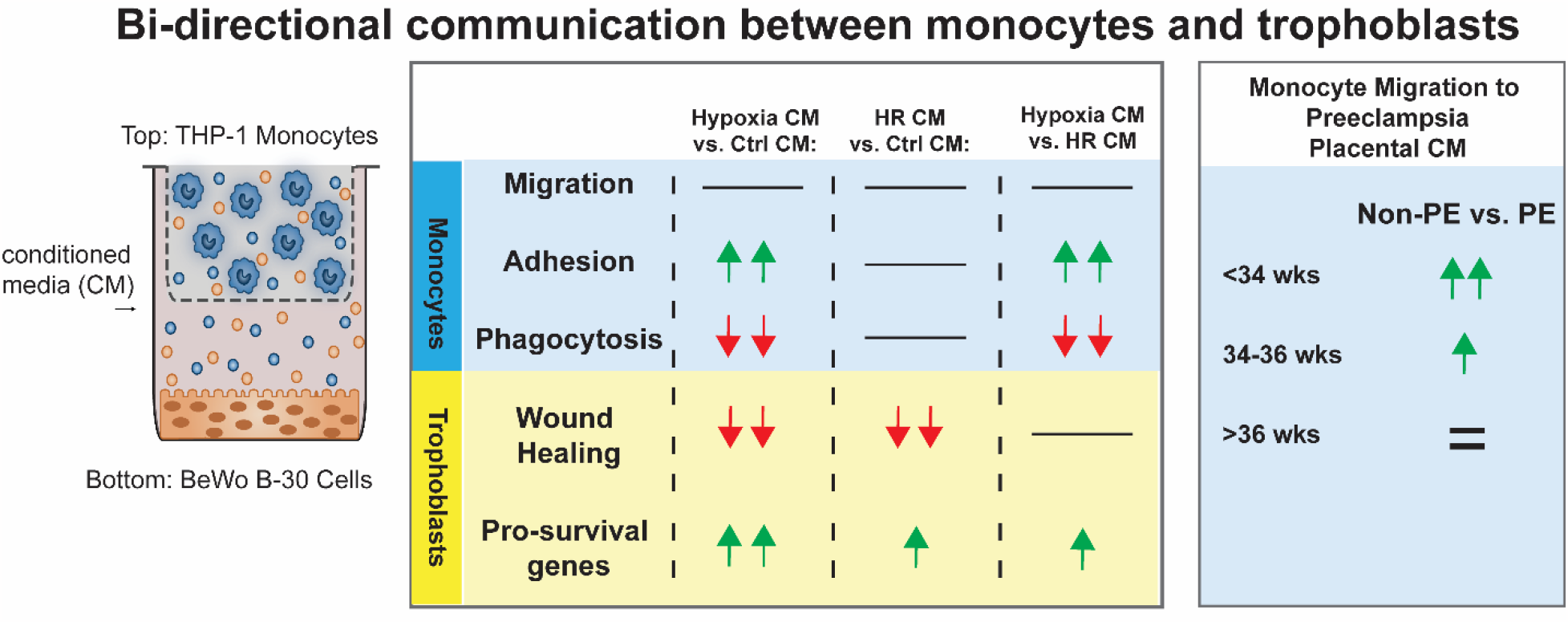

## Introduction

Successful pregnancies require a complex and dynamic relationship between maternal monocytes and placental cells^1^. Monocytes must simultaneously maintain immune protection while providing immunotolerance to the fetus^2–4^. Much of this behavioral adaptation occurs from the influence of placental cells on monocytes as they circulate through the maternal-fetal interface and interact with the syncytiotrophoblast and its molecular signals^5–9^.

Hypertensive disorders of pregnancy (including preeclampsia (PE), 2-8% of pregnancies) are a leading cause of maternal and neonatal mortality^10–12^. Preeclampsia is accompanied by proteinuria or maternal organ dysfunction^13,14^. Current understanding of disease etiology is limited, but data suggests placental malperfusion originates from the stress and damage of the syncytiotrophoblast^15^. This cellular injury alters the population numbers^9^ and bioactivity^16(p13),17^ of the syncytiotrophoblast extracellular vesicles, as well as the production of cytokines^18^ and steroidal hormones^19^. Accordingly, monocytes in preeclampsia patients have altered activity with evidence that they are more pro-inflammatory^3,4,20^ and adhesive^21,22^, but with altered phagocytic^23^ and migrational behavior^22^.

A debate exists in scientific literature whether this placental stress results from prolonged placental hypoxia or a hypoxia-reperfusion (HR) type injury. *In vitro* research comparing the two models of syncytiotrophoblast stress outlines the response of the placental cell itself^24–26^. However, because preeclampsia involves both a maternal factor and a fetal factor, an effective *in vitro* disease model should replicate the interactions observed between the syncytiotrophoblast and maternal monocytes.

In this study, we demonstrate the ability to recapitulate some features of preeclampsia using cell line models of syncytiotrophoblast-monocyte interactions and exposing them to hypoxic signals.

## Methods

### Cell Culture

The choriocarcinoma cell line BeWo b30 was a gift from the Whitehead lab at Carnegie Mellon University. The cells were cultured in F-12k media (ATCC) containing 10% FBS. The monocyte cell line THP-1 (ATCC) was cultured in RPMI 1640 media (ATCC) with 10% FBS.

BeWo b30 cells were cultured in 24-well plates until 70-80% confluent. Cellular hypoxia was induced by treating the cells with serum-free F-12k media containing 300 μM cobalt chloride hexahydrate (CoCl_2_) for 24 hours. The media was then either replaced with CoCl_2_ media again (hypoxic group) or with normal growth media (HR group). For coculture studies, transwell inserts (Sterlitech) with 0.4 μm pores were added to the well post media change. 250 μL of THP-1 monocytes were seeded in the inserts at 2x10^6^ or 4x10^6^ million cells/mL. The coculture system was incubated for 24 hours before experimentation. As a control, monocytes were cultured in the transwell system with CoCl_2_ media.

### Placental Explant Conditioned Media

Placental tissue was obtained through the Steven C. Caritis Magee Obstetric Maternal and Infant Biobank. Limited patient demographics are demonstrated in **(Supplemental Table 1)**. Full thickness placental biopsies were taken from cesarean sections in the absence of labor or ruptured membranes. Tissue was dissected within 2 hours of delivery and maternal decidua was removed. Chorionic villi was dissected off of fetal vessels and 0.5cm x0.5cm segments were placed in 1mL of DMEM supplemented with penicillin-streptomycin and 10% fetal bovine serum. These were incubated at 37 degrees with 5% CO_2_ as previously described. Following incubation, the supernatants were collected and stored at 4 degrees Celsius until further analysis. *qPCR*

RNA was extracted from cells using the miRNAeasy Plus kit (Qiagen) and stored at -80 degrees Celsius until further experimentation. The RNA was converted to cDNA using the high capacity cDNA conversion kit (Invitrogen). qPCR was performed using the following primers (Integrated DNA Technologies): Actin-β, erythropoietin (*EPO*), glucose transporter-1 (*GLUT-1*), vascular endothelial growth factor (*VEGF*), binding of immunoglobulin protein (*BiP*), CCAAT/enhancer-binding protein homologous protein (*Chop*), BcL2-associated X-protein (*BAX*), *BcL2*, microtubule-associated protein 1A/1B-light chain 3 (*LC3*), mammalian target of rapamycin (*mTOR*), *CX3CL1*, and *ICAM-1*. The resulting data was analyzed via the ΔΔCt method and compared to *ACTB*. Monocyte polarization was analyzed via the following primers (Integrated DNA Technologies): *CD64, CD40, CD33, CD86, CD163, CD206, CXCR1, CXCR2, CCR5, HLA-DR*α, *HLA-DR*β*1, HLA-DR*β*3, CCR2, CX3CR1, LFA-1, IL-8, IL-10, IL-12, TNF-*α, *TGF-*β*1, MIP-1*α, *and NOS2*. Primer sequences can be found in **(Supplemental Table 2)**.

### Wound Healing Assay

BeWo b30 cells were cultured in 24-well plates until confluent. A 10 μL pipette tip was used to make vertical scratches in the center of each well. Each scratch was imaged using a Keyence microscope (2x magnification). The cells were then treated with hypoxia as described previously. 0.4 μm transwell inserts containing 5x10^5^ THP-1 monocytes were placed in the well and the system was incubated for 48 hrs. At 24 hrs, the HR group was reperfused. The wells were imaged again after the 48 hour incubation, and wound area was calculated using ImageJ.

### Monocyte Adhesion Assay

The adhesion assay protocol from Siwetz et al was adapted for this study^6^. Briefly, BeWo b30 cells were cultured in 6-well plates until confluency. The three experimental groups were as follows: control (normal growth media), hypoxic (CoCl_2_ treatments for 24 hrs), and the hypoxia-reperfusion group (24 hrs of CoCl_2_ treatment followed by 24 hrs of reperfusion with normal media). 6x10^5^ THP-1 cells were added to each well of placental cells followed by 90 minutes incubation. The adherent cells were washed three times with phosphate-buffered saline and trypsinized to detach the cells. The cell suspension underwent two centrifuge rinses with cell staining buffer, followed by blocking with Trustain FcX solution (Biolegend) and staining with anti-human CD33 antibody conjugated to Brilliant Violet 605 (BV605) (Biolegend). The stained cells were analyzed via flow cytometry.

### Monocyte Phagocytosis Assay

THP-1 monocytes were cocultured with BeWo b30 cells as outlined above. In the final 3 hrs of coculture, 4 μL of red fluorescent heat-treated *E. Coli* (Abcam) was added to each insert. After incubation, the phagocytosis of *E. Coli* was measured by quantifying the percentage of monocytes exhibiting red fluorescence using flow cytometry.

### Monocyte Migration Assay

BeWo b30 cells were cultured in 12-well plates until mostly confluent. The cells were then incubated with serum-free media alone, media with CoCl_2_ (24 hrs), or CoCl_2_ media followed by 24 hrs in serum-free media. Following incubation, the media was removed and placed into wells of a 24-well plate. 1x10^6^ THP-1 cells were seeded into 8 μm transwell inserts (Sterlitech) placed in each well. The cells were allowed to migrate for 22 hrs. After the incubation time, cells in the bottom well were counted.

### Statistics

All statistics were calculated using ANOVA tests.

## Results

### HIF-1α expression positively correlates with CoCl_2_ concentration

To mimic and maintain a hypoxic state, cells were incubated with CoCl_2_ **(Figure 1A)**^27^. CoCl_2_ is an effective chemical inducer of hypoxia-inducible factor-1alpha (HIF-1α)^28^, the master regulator of hypoxia-driven pathways within the cell^29^. The optimal concentration of CoCl_2_ was tested by assessing changes in hypoxia related gene expression in both BeWo b30 cell lysate **(Figure 1B)** and resulting conditioned media **(Figure 1C)**. As previously demonstrated, expression of *HIF-1*α positively correlated with CoCl_2_ concentration^30^ **(Figure 1B)**. Accordingly, a concentration of 300 μM CoCl_2_ was selected for all future experiments.

**Figure 1:**
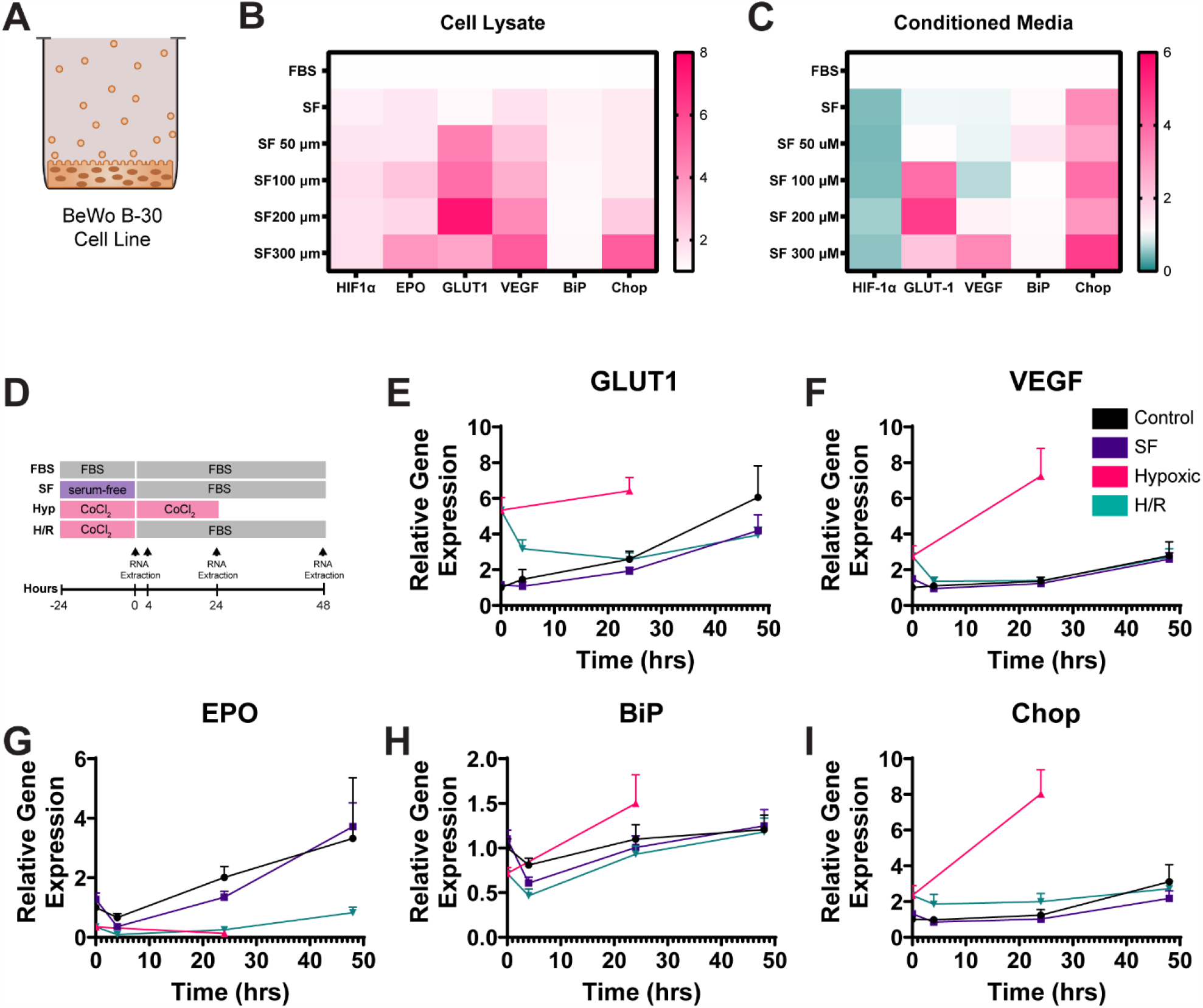
CoCl_2_-induced-hypoxia increases cellular stress while reperfusion decreases stress signals. A) b30 trophoblast cells are incubated with various concentrations of CoCl_2_ for 24 hours. B) and C) Following incubation b30 cell lysate and the conditioned media (CM) were assessed via qPCR for mRNA expression of hypoxia and cell stress related genes (N=2-3 samples for (B), 3 replicates per sample) (N=1-2 samples for (C), 3 replicates per sample). D) b30 cells were treated with either FBS, Serum free media (SF), 300 μm CoCl_2_ (Hyp) or 300μm CoCl_2_ followed by FBS (H/R) to stimulate hypoxia reperfusion. Following media exposure, cell lysate was collected at the varying timepoints (hrs: 0,4,24, and 48) and mRNA expression was measured for the following genes D) Glut1, E) VEGF, F) EPO, G) BiP, and H) Chop. Note: the initial point for both hypoxic and H/R samples is the same sample (N=3 samples per group, N=3 replicates per sample).

### Reperfusion of b30 cells post-hypoxia restores cells to a normoxic state

As HIF-1α expression dissipates quickly upon oxygenation^31^, measuring indirect hypoxia gene expression produces greater reliability in measurement. Thus, we explored differences between hypoxia related gene expression between normoxic, hypoxia, and hypoxia-reperfusion states. Hypoxia-reperfusion was established by incubation with CoCl_2_ for 24 hrs followed by reperfusion with normal growth media over 48 hrs **(Figure 1D)**. We performed qPCR for several genes indirectly induced by hypoxia^32,33^: *EPO, GLUT-1*, and *VEGF*. Hypoxia dramatically increased *VEGF* and *GLUT-1*. However, while VEGF returned to normoxic levels within 4 hrs of reperfusion, *GLUT-1* remained elevated for another 24hrs **(Figure 1E and 1F)**. Endoplasmic reticulum (ER) stress increases the expression of *BiP* and *CHOP. CHOP* remained elevated until 24 hrs post-reperfusion **(Figure 1H-I)**. *BiP* remained lower than the normoxic controls but recovered by 48hrs post-reperfusion **(1H)**. This suggests reperfusion can reduce hypoxia-related cellular stress in BeWo cells but that effects are not immediate.

*EPO* expression differed from the other hypoxia-induced genes **(Figure 1G)** by slowly increasing but never recovering post hypoxia. This contradicted other data confirming both hypoxia and cobalt upregulate EPO^34,35^; however, it confirms a study by EN and Syu demonstrating CoCl_2_ downregulated *EPO* expression in BeWo b30 cells, citing the reason for this unexpected result to be unknown placenta-specific factors^36^.

### Monocytes alter expression of cell death and adhesion genes in hypoxic BeWo b30 Cells

Monocytes respond to hypoxia signals in other tissues^37^. As such, we explored whether monocytes can alter the impact of hypoxia and hypoxia-reperfusion on BeWo b30 cells. Both injury types significantly increased expression of *BAX* (pro-apoptotic) in the BeWo b30 cells **(Figure 2B)**. When monocytes were cocultured with the BeWo cells, they decreased *BAX* expression in the hypoxia group but in neither of the other groups. *BcL2* (pro-survival) increased in hypoxic conditions but decreased in hypoxia-reperfused cells. Monocyte addition elevated *BcL2* in the hypoxia and hypoxia-reperfusion groups, but not in the control. Taken together, **(Figures 2B-C)** demonstrate that monocytes promote the survival of stressed placental cells.

**Figure 2:**
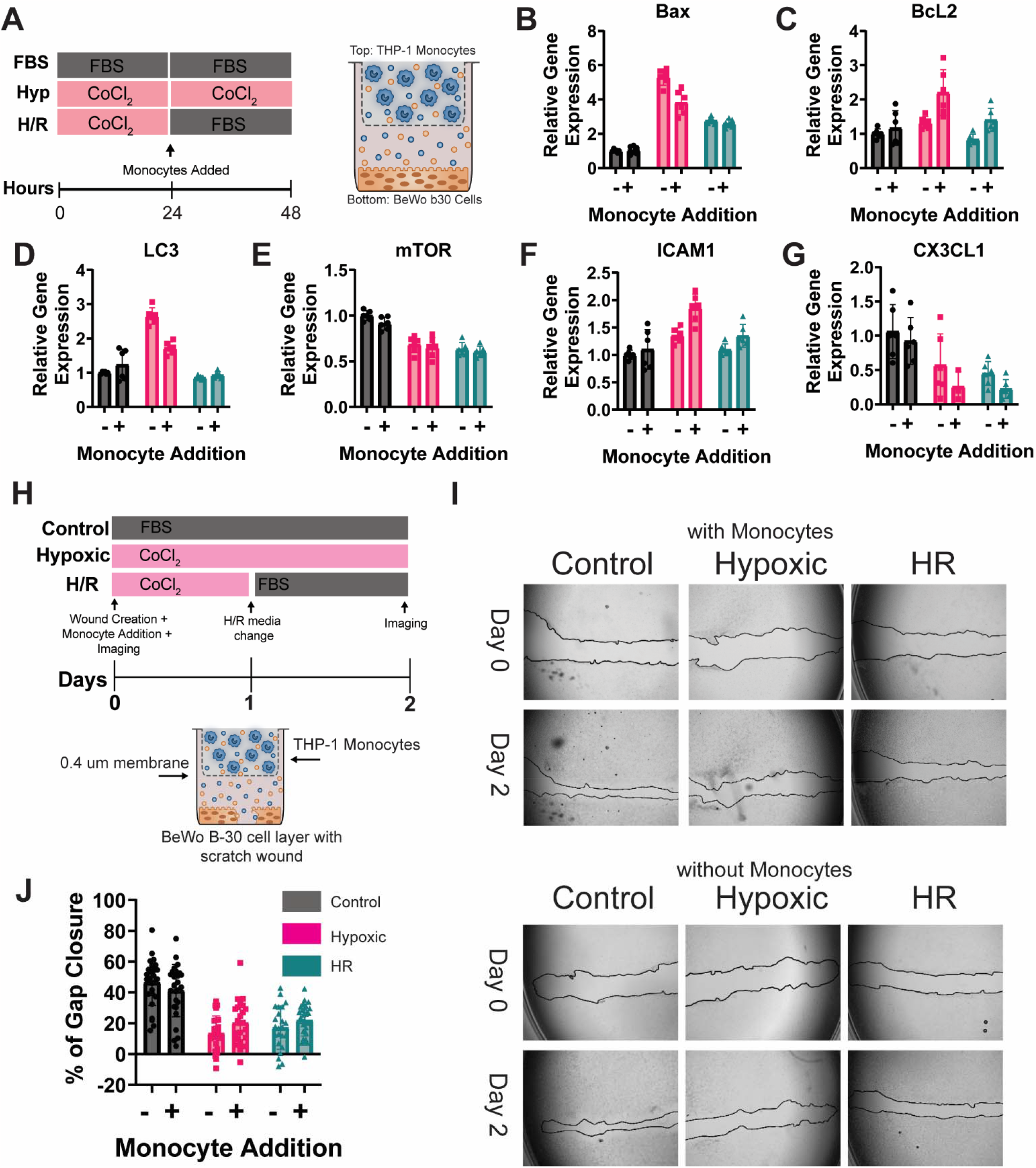
Addition of Monocytes increased cell survival and adhesion gene expression in b30 cells under hypoxia conditions. A) b30 and THP-1 monocytes are incubated in a transwell co-culture with an 0.4μm filter under FBS, Hypoxic (Hyp), or Hypoxic Reperfusion conditions (H/R). b30 mRNA is measured after 48hrs of culture without monocytes or co-incubation with monocytes for the following genes: B) BAX, C) BcL2, D) LC3, E) mTOR, F) ICAM-1, and G) CX3CL1(N=2 cocultures per group, N=3 replicates per coculture). Results were analyzed via a two-way ANOVA. Placental State: p<0.0001 for all genes. Monocyte Presence: p=0.0033 (BAX), p<0.0001 (BcL2), p=0.0094 (LC3), p=0.0633 (mTOR), p<0.0001 (ICAM1), p=0.0268 (CX3CL1). Interaction: p<0.0001 (BAX and LC3), p=0.0004 (BcL2), p=0.5763 (mTOR), p=0.0058 (ICAM1), p=0.3821 (CX3CL1). H) Using the same transwell setup, a wound healing scratch is induced on the b30 monoculture layer. I) Representative brightfield images of the wound healing area Day 0 and 2 after the scratch is initiated. Magnification = 2X (Keyence microscope) J) Quantifications of the gap closure (percentage) at the end of 48hrs calculated from the images. Statistics were calculated using a two-way ANOVA (J). Placental State: p<0.0001, Monocyte Presence: p=0.3418, Interaction Factor: p=0.0563 (N=23-28 scratches per group).

The genes *LC3* and *mTOR* correlate with cellular autophagy. **(Figure 2D)** demonstrates hypoxia increases the expression of *LC3* (an autophagosome marker) and monocyte addition reduces the magnitude of hypoxia induced expression (p<0.01). This suggests monocytes exert a protective effect against autophagy in hypoxic cells. *mTOR* inhibits autophagy and was decreased in both hypoxic and hypoxic-reperfused cells **(Figure 2E)**, but monocyte addition did not alter its expression. Consequently, these results suggest monocytes have a selective, specific function in regulating genes involved in autophagy.

Next, we examined the effect hypoxia and hypoxia-reperfusion had on adhesion genes. Hypoxia alone elevated *ICAM-1* expression in placental cells **(Figure 2F)**. In contrast, hypoxia-reperfusion alone significantly decreased *CX3CL1* **(Figure 2G)**. The presence of monocytes further upregulated *ICAM-1* and downregulated *CX3CL1* expression in both the hypoxic and hypoxic-reperfused groups. In the control, monocyte presence did not change *ICAM-1* expression and only slightly decreased the expression of *CX3CL1*. These results imply that under conditions of oxygen deprivation, monocyte adhesion to the STB is primarily facilitated via the ICAM-1 ligand.

### Placental injury decreases wound healing

As preeclampsia causes damage to the syncytiotrophoblast, we examined the effects of placental injury and monocyte presence on cellular wound-healing. Both hypoxia and HR injury significantly decreased the wound-healing abilities of the BeWo b30 cells compared to the control. The presence of monocytes did not influence the wound-healing capabilities of trophoblasts in any group **(Figure 2 I-J)**.

### Selection of a panel of monocyte polarization genes

Monocyte polarization refers to the spectrum of monocyte phenotypes and behaviors associated with varying degrees of inflammation^38^. Broadly, this spectrum is stratified into three subpopulations based on monocyte expression of surface receptors CD14 and CD16. Classical monocytes (CD14++CD16-) are professional phagocytes (80-95%), non-classical monocytes (CD14+CD16+) patrol the endothelium (2-11%), and intermediate monocytes (CD14++CD16+) comprise the remainder^39^. We selected a panel of surface receptors and cytokines that are differentially expressed amongst the three subpopulations **(Supplemental Figure 1)**.

### Placental state influences monocyte polarization

Numerous studies have attempted to quantify shifts in monocyte subpopulations in preeclampsia. The results conflict whether non-classical or intermediate populations are most upregulated, but it is generally accepted that CD16-monocytes decrease and CD16+ monocyte populations increase^3,4,20,40,41^, similar to autoimmune diseases^42^.

Therefore, we assessed whether signals from hypoxic or hypoxic-reperfused BeWo b30 cells could influence monocyte polarization **(Figure 3A)**. Our results show placental hypoxia increased the gene expression of several genes associated with the CD16+ monocytes. Placental hypoxia increased the expression of *CD163, CD86, CD40, HLA-DR*α, *IL-8, TNF-*α, *TGF-*β*1*, and *MIP-1*α compared to all other groups **(Figure 3B)**. *HLA-DR*β*1* was increased in the hypoxic group relative to the control group alone. Of these genes, HR increased *CX3CR1, CXCR2, CCR2, CD86, HLA-DR*β*3, TNF-*α, and *TGF-*β*1* but decreased *MIP-1*α expression relative to control samples. *IL-8* expression in HR was decreased relative to all groups **(Figure 3B)**.

**Figure 3:**
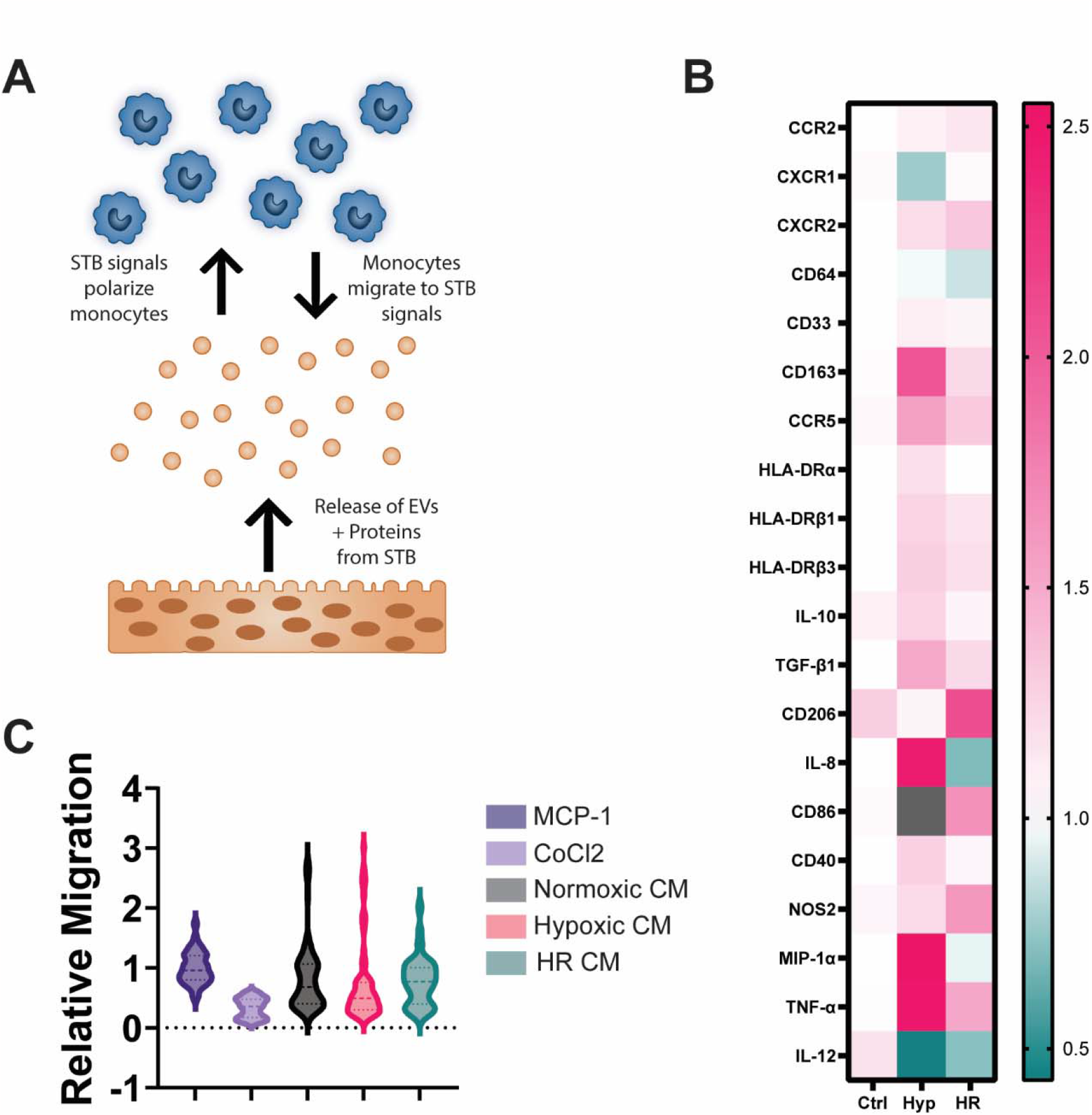
Changes in THP-1 monocyte mRNA expression and chemotaxis following exposure to hypoxia and hypoxia reperfusion. (A) STB signaling affects monocyte polarization and migration. (B) THP-1 monocytes were cocultured with BeWo b30 cells using a transwell coculture system with 0.4 μm pores. After 24 hrs of coculture, RNA was extracted from the monocytes in the top well and qPCR analysis determined the expression of various genes pertaining to monocyte polarization (N=2 cocultures per group, N=1-3 replicates per coculture) (C) BeWo b30 conditioned media was placed in the bottom well of a transwell migration chamber and naïve THP-1 monocytes were placed in the top insert (8 μm pores). Cellular migration was quantified relative to monocyte migration to MCP-1. Statistics were calculated using a one-way ANOVA. All comparisons between groups were not significant (p>0.05) except for the pairs involving the CoCl_2_ control (N=4 transwells per group, N=4-7 replicates per transwell).

As a control, THP-1 monocytes were cultured in the transwell system with CoCl_2_, which altered the expression of 18 of the 22 genes compared with untreated monocytes. CoCl_2_ upregulated *CD86, CCR5, CXCR1, CD163*, all *HLA-DR* sequences, *CD40, IL-8, IL-10, TNF-*α, *TGF-*β*1, NOS2*, and *MIP-1*α. Expression of *CXCR2* and *IL-12* was decreased following CoCl_2_ treatment **(Supplemental Figure 1)**.

### Monocyte migration to BeWo b30 conditioned media

CXCR1, CXCR2, and CCR2 are receptors for monocyte chemokines, with IL-8 binding to the first two, and MCP-1 binding to CCR2. qPCR analysis revealed *CXCR1* and *CCR2* expression in monocytes did not change between any groups. *CXCR2* gene expression was significantly increased in the HR group vs. the normoxic group. **(Figure 3B)**.

As such, we examined whether signals from hypoxic and hypoxic-reperfused BeWo b30 cells induce monocyte migration. Monocyte chemoattractant protein-1 (MCP-1) was used as a positive control for migration. When plotted as a ratio to MCP-1 migration, CoCl_2_ significantly decreased migration compared to all groups. No significant differences were observed in the coculture groups vs. the MCP-1 control or between the normoxic, hypoxic, and HR groups **(Figure 3C)**. These results imply CoCl_2_ suppresses migration, but placental stress does not impact monocyte chemotaxis.

### Placental hypoxia decreases monocyte phagocytosis

As “first responders” of the immune system, monocytes are responsible for phagocytosing pathogens. Consequently, we examined how signals from injured BeWo b30 cells impact THP-1 phagocytosis of *E. Coli*. Cytochalasin D inhibits phagocytosis and was used as a negative control^43^. Cytochalasin D slightly decreased phagocytosis compared to untreated controls; however, the results were not significant. Culturing THP-1 cells with normoxic or HR BeWo b30 cells significantly increased their phagocytosis by a factor of <5x compared to the other experimental groups (p<0.0001). However, this increase in phagocytosis was not observed in THP-1 cells cultured with hypoxic placental cells **(Figure 4)**.

**Figure 4:**
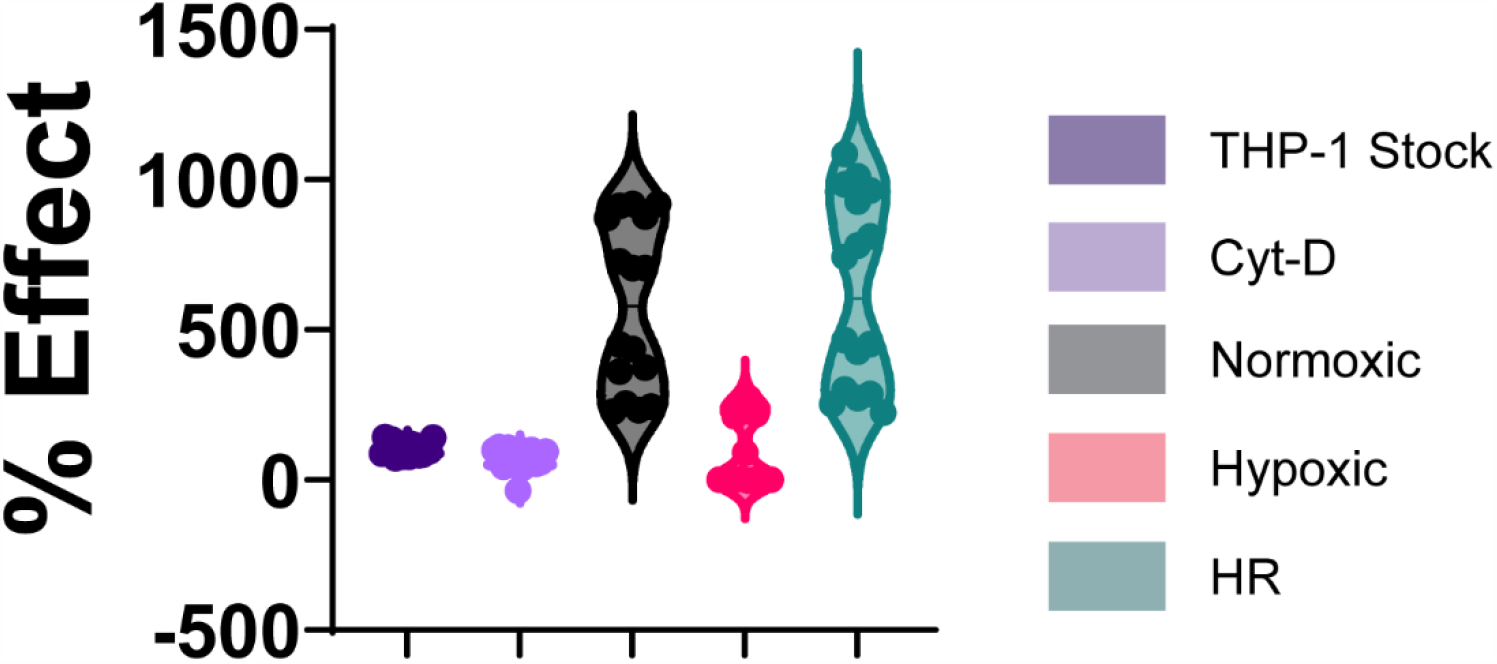
THP-1 monocytes exhibit decreased phagocytosis under hypoxic conditions. THP-1 monocytes were cocultured with BeWo b30 cells for 24 hrs using the transwell coculture system. In the last 4 hrs of incubation, red fluorescent E. Coli were added to each insert. THP-1 internalization of fluorescent E. Coli was assessed via flow cytometry. Results were expressed as a % effect compared to the phagocytosis of the untreated THP-1 cells. Statistics were calculated using a one-way ANOVA. All groups with significance had a p value of <0.0001 (N=7-8 cocultures per group, N=2 replicates per coculture).

### Placental hypoxia decreases monocyte adhesion

Monocytes circulating through the intervillous space can adhere to the syncytiotrophoblast. Previous research demonstrated extracellular vesicles from preeclampsia patients increased the adhesiveness of THP-1 monocytes^22^. We assessed whether signals derived from BeWo b30 cells could affect the direct adhesion of monocytes to their surface. Our results demonstrated THP-1 cells adhered to placental cells at a higher rate under hypoxia compared with other experimental groups **(Figure 5B)**.

**Figure 5:**
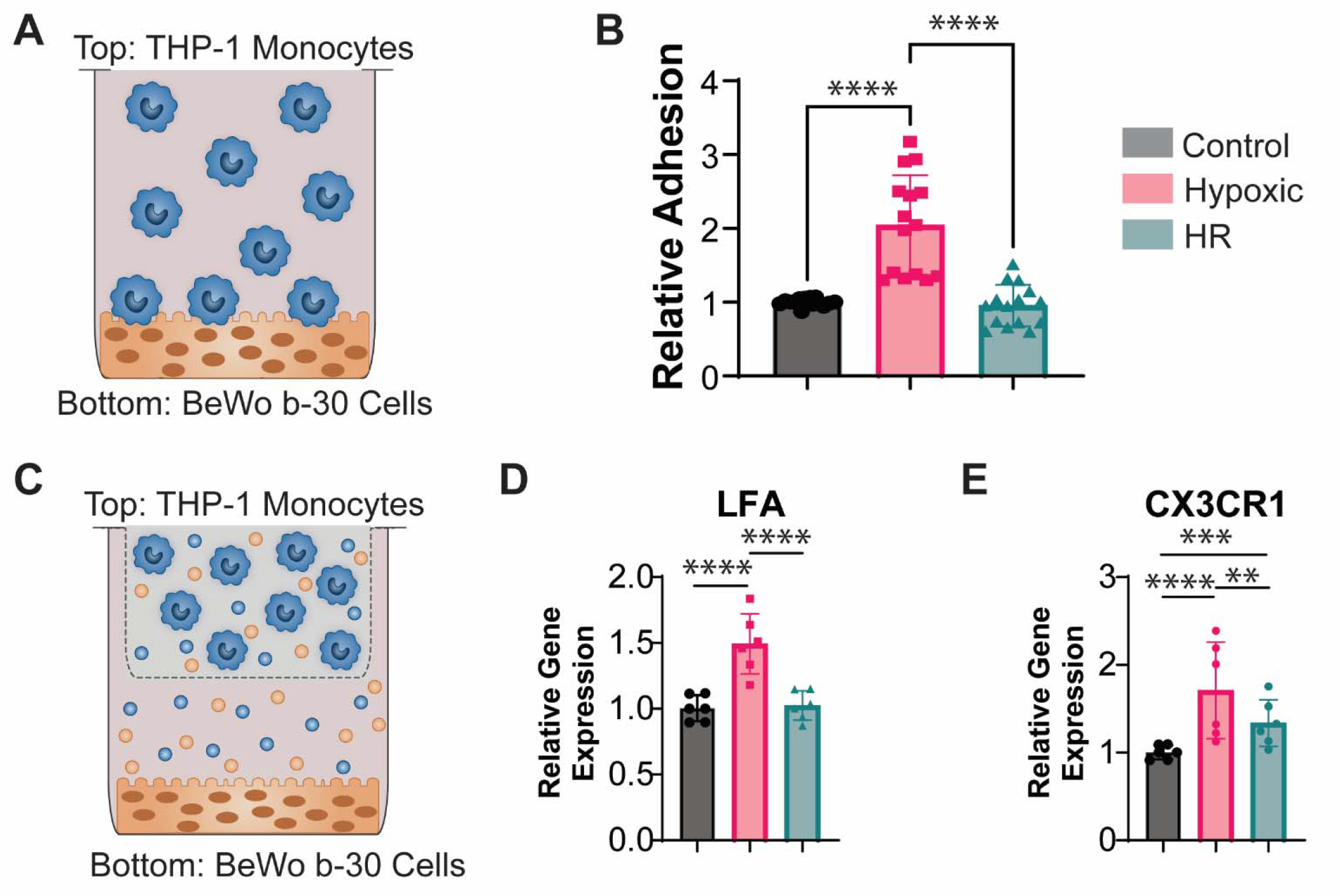
Adhesion is increased under hypoxia. Naïve THP-1 monocytes were added to cultured BeWo b30 cells and allowed to incubate for 90 minutes (A). The number of adherent monocytes was determined by flow cytometry. Results were expressed relative to THP-1 adhesion to normoxic BeWo b30 cells. Statistics were calculated via a one-way ANOVA. All groups had a p value of <0.0001 compared to the hypoxic group (N=6 cocultures per group, N=2-3 replicates per coculture) (B). THP-1 cells were cocultured with monocytes using a transwell coculture system with 0.4 μm pores (C). After 24 hrs of coculture, RNA was extracted from the monocytes and qPCR was performed to determine the expression of monocyte adhesion receptor genes (D-E). Statistics were calculated using a one-way ANOVA. For LFA, all groups had a p value of <0.0001 compared to the hypoxic group. For CX3CR1: p<0.001 (ctrl vs. hypoxic), p=0.0006 (ctrl vs. HR), p=0.0042 (hypoxic vs. HR) (N=2 cocultures per group, N=3 replicates per coculture).

Furthermore, we assessed the expression of monocyte adhesion receptor genes in THP-1s following coculture with BeWos **(Figure 5C)**. Monocytes bind to the ICAM-1 ligand via the LFA-1 receptor^44^ and to CX3CL1 via the CX3CR1 receptor^6^. The presence of placental hypoxia but not HR increased *LFA-1* and *CX3CR1* gene expression compared to the normoxic coculture **(Figure 5 D-E)**. As a control, monocytes were cultured in the transwell assay system with CoCl_2_. CoCl_2_ significantly increased the expression of both *LFA-1* and *CX3CR1* **(Supplemental Figure 1)**.

### Early-Onset Preeclampsia impacts THP-1 monocyte migration to placental explant signals

As limited research exists regarding monocyte chemotaxis to STB signaling, we analyzed villous placental explant conditioned media for the presence of chemokine RNA as well as its ability to induce monocyte chemotaxis. RNA isolated from placental explant conditioned media revealed the presence of monocyte chemokine genes *MCP-1, CX3CL1*, and *IL-8* across various gestational ages **(Figure 6 A,B, D)**.

**Figure 6:**
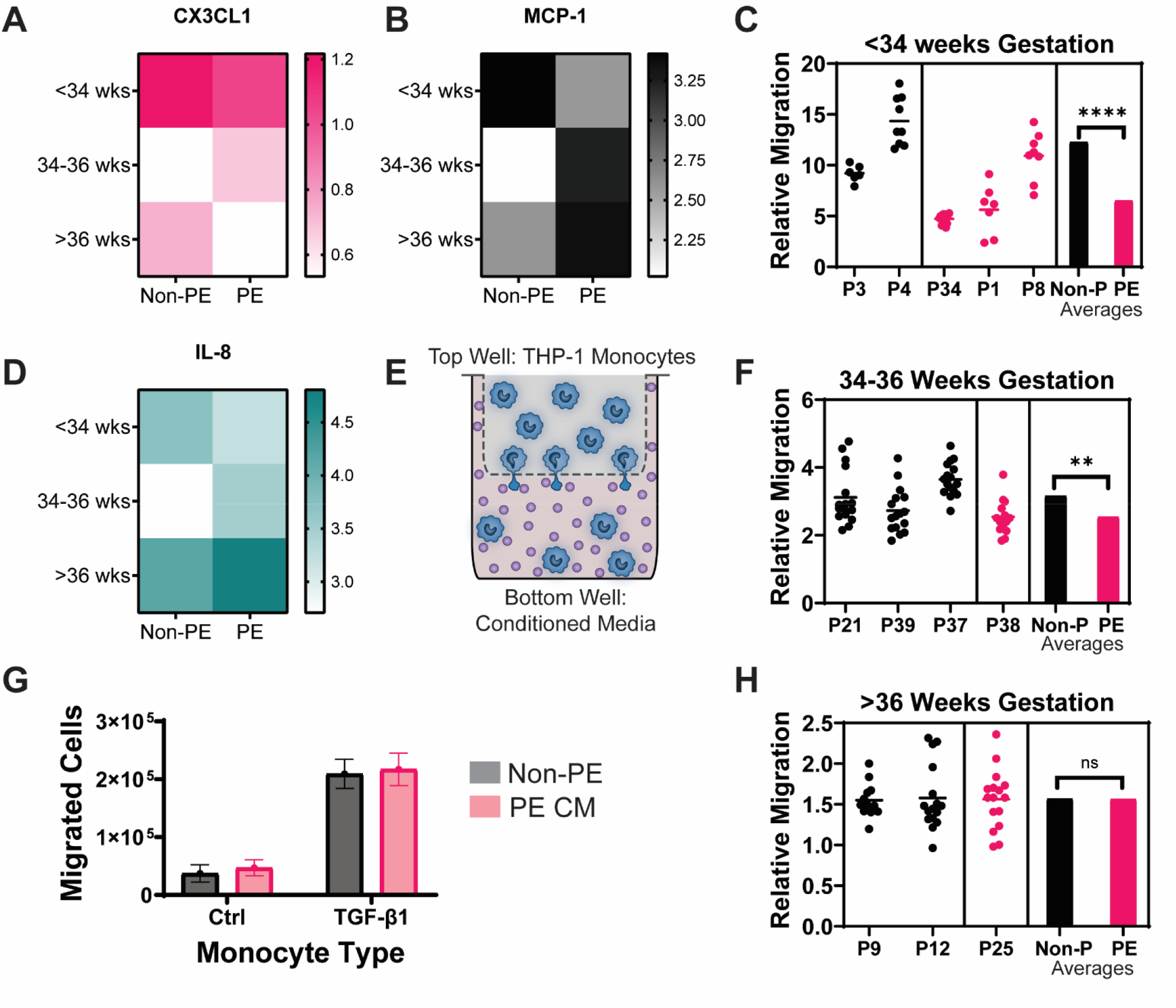
Monocyte migration to human placental explant conditioned media. Placental explants were obtained from caesarean deliveries at Magee-Womens hospital. Villous tissue was dissected from the explant and incubated in media for 24 hours. RNA was extracted from the placental explant conditioned media and qPCR analysis revealed the presence of monocyte chemokine RNA (A, B, and D) (N=3 replicates per placenta). The placental explant media was placed into the bottom well of a transwell migration chamber and THP-1 monocytes were placed in the top insert (E). Monocyte migration relative to 50 ng/mL MCP-1 was quantified across various gestational ages and results were analyzed using a one-way ANOVA (C, F, H) p<0.0001 for (C), p=0.0032 for (F), and p=0.9713 for (H) (N=3-4 transwell chambers per placenta). The transwell migration assay was repeated using naïve THP-1 monocytes and those treated with 10 ng/mL TGF-β1 with placental conditioned media between 34 and 35 weeks gestation (G) (N=1-2 placentas per disease group, N=4 transwell chambers per placenta). The statistics were calculated using a two-way ANOVA. Placental state: p=0.0574, monocyte treatment: p<0.0001, interaction: p=0.8230.

THP-1 monocyte migration to placental explant conditioned media strongly depended upon the gestational age of the placenta **(Figure 6 C, F, H)**. For samples <34 weeks gestation, migration was significantly decreased (p<0.0001) in PE vs. the non-PE placentas **(Figure 6C)**. This trend continued in the 34-36 week age group, but not as significantly as the youngest group **(Figure 6F)**. Finally, for samples >36 weeks gestation, monocytes migrated equally to non-PE and PE explant media **(Figure 6H)**.

Since monocyte subpopulations shift towards the CD16+ populations in preeclampsia, we explored whether placental disease state or monocyte phenotype most strongly affects monocyte chemotaxis. TGF-β1 is a cytokine which induces CD16 expression on monocytes^45^. When comparing the effects of placental state vs. monocyte condition, it was observed that monocytes preconditioned with TGF-β1 exhibited significantly higher rates of migration compared to untreated THP-1s **(Figure 6G)**. This data implies monocyte state impacts chemotaxis more than placental signaling.

## Discussion

The underlying pathogenesis for hypertensive disorders of pregnancy remains unclear and both hypoxia and hypoxia-reperfusion injury to trophoblasts have been implicated. We undertook this study to determine how these models modulate syncytiotrophoblast-monocyte communication and to elucidate how monocytes respond functionally to syncytiotrophoblast damage.

Our study demonstrates a model of syncytiotrophoblast stress that recapitulates several hallmarks of preeclampsia. *HIF-1*α, the master regulator of hypoxia^29^, is overexpressed in the preeclamptic placenta^26,46^ and in pregnancies at high altitudes^47^, which have an increased risk of developing preeclampsia due to decreased oxygen tension^48,49^. Placental hypoxia can induce *VEGF* production similar to PE in primary human trophoblast cells^26^. Our *in vitro* model confirmed these previous findings by demonstrating only prolonged hypoxia in BeWo b30s upregulated *HIF-1*α and *VEGF*.

Moreover, microscopic evaluation of the preeclamptic syncytiotrophoblast reveals enlarged endoplasmic reticulum, indicative of ER stress^15,50^. Our results demonstrated hypoxia induced ER stress genes *BiP and CHOP*, similar to a previous study indicating hypoxia but not hypoxia-reperfusion injury caused upregulation of ER stress genes^26^.

Placental apoptosis is increased in the preeclamptic placenta^25^. The apoptosis pathway is regulated via the *BcL2* gene family where *BAX* promotes apoptosis and *BcL2* blocks *BAX* oligomerization to inhibit apoptosis^51,52^. Previous research indicated that *BcL2* protein is localized in the syncytiotrophoblast but *BAX* was not expressed^52^. In contrast, our data found that BeWo b30 cells express *BAX*. Our results indicate placental hypoxia and not hypoxia-reperfusion most strongly drives cellular apoptosis via the upregulation of *BAX* and the downregulation of *BcL2*.

Literature remains unclear whether placental autophagy increases or decreases in preeclampsia^53^. LC3 positively regulates autophagosome formation, and Oh et al identified that severe preeclampsia upregulates placental protein expression of LC3^54^. Our data revealed hypoxia mimicked the results obtained by Oh et. al using hypoxic JEG cells^54^. Staining for mTOR in PE placentas has been shown to be decreased compared to healthy controls^56^, while another study found mTOR expression in the syncytiotrophoblast was not significantly different in PE vs. healthy placentas^57^. Our data showed placental injury decreased mTOR expression, indicating similar to the LC3 results that the pro-autophagy pathway was enhanced in hypoxia.

To our knowledge, this study is the first to examine an *in vitro* model of syncytiotrophoblast-monocyte interactions in the context of preeclampsia. Our results demonstrate several critical functions of peripheral monocytes as they interact with the syncytiotrophoblast. First, monocytes mitigated apoptosis in BeWo b30 cells by decreasing *BAX* and increasing *BcL2*, and selectively mitigated part of the autophagy pathway by downregulating *LC3*, implying that monocytes prevent cellular death in the damaged syncytrophoblast.

However, monocyte presence did not affect wound-healing abilities of the injured BeWo b30 cells. As the syncytiotrophoblast is formed via the fusion of the cytotrophoblast cells^58^, placental injury is hypothesized to contribute to “cracks” in the syncytiotrophoblast, allowing fetal hemoglobin to escape into maternal circulation^59,60^. Previous studies remain unclear whether the presence of immune cells improves or further damages these wounds^15^, and our results are consistent with these studies.

Next, when the influence of placental state on monocyte polarization and adhesion was compared, 10 genes investigated were increased in the hypoxic group compared to all other groups: 6 surface receptors (*CX3CR1, CD86, CD163, CD40, HLA-DR*α, *LFA-1*) and 4 cytokines (*IL-8, TNF-*α, *TGF-*β*1, MIP-1*α). Of the genes increased in the hypoxic group, all of them pertain to proteins predominantly expressed by CD16+ monocyte populations^61–65^. Our data implies placental hypoxia most strongly influences monocyte populations toward a CD16+ phenotype.

Circulating monocytes bind to the syncytiotrophoblast via LFA-1/ICAM-1^44^ or CX3CR1/CX3CL1 interactions between the cells^5,6^. Our data revealed that hypoxia increased *ICAM-1* in BeWo b30s and monocyte addition further upregulated this gene. However, it is well established that non-classical monocytes use LFA-1 to patrol the endothelium^66^. Taken together, these results indicate hypoxia primes the STB for increased monocyte adhesion and bi-directional communication between the cells facilitates monocyte binding to the placental surface via LFA-1/ICAM-1 interaction. The reverse trend was observed for placental CX3CL1 expression: both hypoxia and HR decreased gene expression and monocytes further decreased this. This indicates the predominant means by which monocytes bind to the injured STB is via ICAM-1/LFA-1 interactions.

Hypoxic placental cells decreased monocyte phagocytosis, increased monocyte adhesion, and did not change monocyte migration toward placental signals. Monocytes isolated from PE patients exhibit decreased phagocytosis in comparison with monocytes from healthy pregnant women^23^. Monocyte adhesion to the endothelium increases after treatment with PE plasma^21,67^, and plasma-derived microparticles from PE patients have been demonstrated to decrease migration and increase monocyte adhesiveness^22^. When replicating monocyte functional behavior in PE, our results show that hypoxia is a more effective placental model.

Our findings indicate monocyte migration to placental factors secreted in early-onset PE (<34 weeks gestation) is decreased compared to non-PE samples. This implicates the potential differences in etiology between early-onset preeclampsia and late-onset preeclampsia. It remains unanswered whether these migration changes are a protective mechanism or a potential therapeutic target. Our migration results using the *in vitro* models most accurately represent the migration data observed in late-onset PE, implying that additional factors besides oxidative stress may be impacting monocyte chemotaxis.

Finally, monocyte inflammation more strongly influenced migration than placental disease state. This holds potential applications in the development of diagnostic assays by which one could predict monocyte functional behavior at the maternal-fetal interface and the onset of disease.

Overall, this *in vitro* model of preeclampsia addresses two primary gaps of knowledge. First, it analyzes proposed models of preeclampsia by studying cell-cell interactions instead of only surveying placental tissue. Second, many existing studies involving trophoblast-monocyte interactions involve either decidual or fetal macrophages. This model provides a platform to study monocyte-trophoblast interactions within the intervillous space, and how placental disease states affect these interactions.

## Conclusion

The conclusions of this study are threefold: First, CoCl_2_ effectively induces a hypoxic state within placental cells. Second, monocytes promote survival in hypoxic placental cells but do not alter their wound-healing capacity. Third, compared to HR placental cells, hypoxic placental cells induce monocyte phenotype and behavior similar to PE monocytes. Overall, our results indicate that placental hypoxia more accurately recapitulates the bi-directional relationship between monocytes and the syncytiotrophoblast in preeclampsia.

## Acknowledgments

The authors thank Dr. Katie Whitehead for the BeWo b30 cells. This work was funded (in part) by the Dowd Fellowship from the College of Engineering at Carnegie Mellon University. The authors would like to thank Philip and Marsha Dowd for their financial support and encouragement.

## Ethics Statement

The authors confirm this article adheres to all the ethical policies of this journal. No ethical approval was required as all research in this article was conducted on commercial cell lines.

## CrediT Author Statement

**Hannah Yankello:** conceptualization, methodology, validation, formal analysis, investigation, writing-original draft, writing-review & editing, visualization **Yerim Lee:** investigation and formal analysis **Dr. Christina Megli:** conceptualization, validation, resources, writing-review & editing, supervision **Dr. Elizabeth Wayne:** conceptualization, methodology, validation, writing-review & editing, visualization, supervision, project administration, funding acquisition.

## Notes

### Competing Interest Statement

The authors have declared no competing interest.

## References

1. Gomez-Lopez N, Guilbert LJ, Olson DM. Invasion of the leukocytes into the fetal-maternal interface during pregnancy. J Leukoc Biol. 2010;88(4):625–633. doi:10.1189/jlb.1209796

2. Veenstra van Nieuwenhoven AL, Heineman MJ, Faas MM. The immunology of successful pregnancy. Hum Reprod Update. 2003;9(4):347–357. doi:10.1093/humupd/dmg026

3. Al-ofi E, Coffelt SB, Anumba DO. Monocyte Subpopulations from Pre-Eclamptic Patients Are Abnormally Skewed and Exhibit Exaggerated Responses to Toll-Like Receptor Ligands. PLOS ONE. 2012;7(7):e42217. doi:10.1371/journal.pone.0042217

4. Melgert BN, Spaans F, Borghuis T, et al. Pregnancy and Preeclampsia Affect Monocyte Subsets in Humans and Rats. PLoS ONE. 2012;7(9):e45229. doi:10.1371/journal.pone.0045229

5. Nonn O, Güttler J, Forstner D, et al. Placental CX3CL1 is Deregulated by Angiotensin II and Contributes to a Pro-Inflammatory Trophoblast-Monocyte Interaction. Int J Mol Sci. 2019;20(3):E641. doi:10.3390/ijms20030641

6. Siwetz M, Sundl M, Kolb D, et al. Placental fractalkine mediates adhesion of THP-1 monocytes to villous trophoblast. Histochem Cell Biol. 2015;143(6):565–574. doi:10.1007/s00418-014-1304-0

7. Göhner C, Fledderus J, Fitzgerald JS, et al. Syncytiotrophoblast exosomes guide monocyte maturation and activation of monocytes and granulocytes. Placenta. 2015;9(36):A47–A48. doi:10.1016/j.placenta.2015.07.329

8. Sacks GP, Clover LM, Bainbridge DR, Redman CW, Sargent IL. Flow cytometric measurement of intracellular Th1 and Th2 cytokine production by human villous and extravillous cytotrophoblast. Placenta. 2001;22(6):550–559. doi:10.1053/plac.2001.0686

9. Germain SJ, Sacks GP, Sooranna SR, Soorana SR, Sargent IL, Redman CW. Systemic inflammatory priming in normal pregnancy and preeclampsia: the role of circulating syncytiotrophoblast microparticles. J Immunol Baltim Md 1950. 2007;178(9):5949–5956. doi:10.4049/jimmunol.178.9.5949

10. Ives CW, Sinkey R, Rajapreyar I, Tita ATN, Oparil S. Preeclampsia-Pathophysiology and Clinical Presentations: JACC State-of-the-Art Review. J Am Coll Cardiol. 2020;76(14):1690–1702. doi:10.1016/j.jacc.2020.08.014

11. Goel A, Maski MR, Bajracharya S, et al. Epidemiology and Mechanisms of De Novo and Persistent Hypertension in the Postpartum Period. Circulation. 2015;132(18):1726–1733. doi:10.1161/CIRCULATIONAHA.115.015721

12. Magee LA, Pels A, Helewa M, Rey E, von Dadelszen P. Diagnosis, evaluation, and management of the hypertensive disorders of pregnancy. Pregnancy Hypertens Int J Womens Cardiovasc Health. 2014;4(2):105–145. doi:10.1016/j.preghy.2014.01.003

13. ACOG Practice Bulletin No. 202: Gestational Hypertension and Preeclampsia. Obstet Gynecol. 2019;133(1):1. doi:10.1097/AOG.0000000000003018

14. Brown MA, Magee LA, Kenny LC, et al. Hypertensive Disorders of Pregnancy: ISSHP Classification, Diagnosis, and Management Recommendations for International Practice. Hypertens Dallas Tex 1979. 2018;72(1):24–43. doi:10.1161/HYPERTENSIONAHA.117.10803

15. Redman CWG, Staff AC, Roberts JM. Syncytiotrophoblast stress in preeclampsia: the convergence point for multiple pathways. Am J Obstet Gynecol. 2021;0(0). doi:10.1016/j.ajog.2020.09.047

16. Sammar M, Dragovic R, Meiri H, et al. Reduced placental protein 13 (PP13) in placental derived syncytiotrophoblast extracellular vesicles in preeclampsia - A novel tool to study the impaired cargo transmission of the placenta to the maternal organs. Placenta. 2018;66:17–25. doi:10.1016/j.placenta.2018.04.013

17. Tannetta DS, Dragovic RA, Gardiner C, Redman CW, Sargent IL. Characterisation of Syncytiotrophoblast Vesicles in Normal Pregnancy and Pre-Eclampsia: Expression of Flt-1 and Endoglin. PLoS ONE. 2013;8(2):e56754. doi:10.1371/journal.pone.0056754

18. Szarka A, Rigó J, Lázár L, Bekő G, Molvarec A. Circulating cytokines, chemokines and adhesion molecules in normal pregnancy and preeclampsia determined by multiplex suspension array. BMC Immunol. 2010;11(1):59. doi:10.1186/1471-2172-11-59

19. Açikgöz ş, Özmen Bayar Ü, Can M, et al. Levels of Oxidized LDL, Estrogens, and Progesterone in Placenta Tissues and Serum Paraoxonase Activity in Preeclampsia. Mediators Inflamm. 2013;2013:862982. doi:10.1155/2013/862982

20. Alahakoon TI, Medbury H, Williams H, Fewings N, Wang XM, Lee VW. Distribution of monocyte subsets and polarization in preeclampsia and intrauterine fetal growth restriction. J Obstet Gynaecol Res. 2018;44(12):2135–2148. doi:10.1111/jog.13770

21. Flood-Nichols SK, Stallings JD, Gotkin JL, Tinnemore D, Napolitano PG, Ippolito DL. Elevated ratio of maternal plasma ApoCIII to ApoCII in preeclampsia. Reprod Sci Thousand Oaks Calif. 2011;18(5):493–502. doi:10.1177/1933719110390390

22. Kovács ÁF, Láng O, Turiák L, et al. The impact of circulating preeclampsia-associated extracellular vesicles on the migratory activity and phenotype of THP-1 monocytic cells. Sci Rep. 2018;8(1):5426. doi:10.1038/s41598-018-23706-7

23. Lampé R, Kövér Á, Szűcs S, et al. Phagocytic index of neutrophil granulocytes and monocytes in healthy and preeclamptic pregnancy. J Reprod Immunol. 2015;107:26–30. doi:10.1016/j.jri.2014.11.001

24. Hung TH, Skepper JN, Burton GJ. In vitro ischemia-reperfusion injury in term human placenta as a model for oxidative stress in pathological pregnancies. Am J Pathol. 2001;159(3):1031–1043. doi:10.1016/S0002-9440(10)61778-6

25. Hung TH, Skepper JN, Charnock-Jones DS, Burton GJ. Hypoxia-reoxygenation: a potent inducer of apoptotic changes in the human placenta and possible etiological factor in preeclampsia. Circ Res. 2002;90(12):1274–1281. doi:10.1161/01.res.0000024411.22110.aa

26. Fuenzalida B, Kallol S, Zaugg J, et al. Primary Human Trophoblasts Mimic the Preeclampsia Phenotype after Acute Hypoxia–Reoxygenation Insult. Cells. 2022;11(12):1898. doi:10.3390/cells11121898

27. Muñoz-Sánchez J, Chánez-Cárdenas ME. The use of cobalt chloride as a chemical hypoxia model. J Appl Toxicol. 2019;39(4):556–570. doi:10.1002/jat.3749

28. Tripathi VK, Subramaniyan SA, Hwang I. Molecular and Cellular Response of Co-cultured Cells toward Cobalt Chloride (CoCl2)-Induced Hypoxia. ACS Omega. 2019;4(25):20882–20893. doi:10.1021/acsomega.9b01474

29. Weidemann A, Johnson RS. Biology of HIF-1α. Cell Death Differ. 2008;15(4):621–627. doi:10.1038/cdd.2008.12

30. Shi J, Wan Y, Di W. Effect of hypoxia and re-oxygenation on cell invasion and adhesion in human ovarian carcinoma cells. Oncol Rep. 2008;20(4):803–807. doi:10.3892/or_00000077

31. Huang LE, Arany Z, Livingston DM, Bunn HF. Activation of Hypoxia-inducible Transcription Factor Depends Primarily upon Redox-sensitive Stabilization of Its α Subunit *. J Biol Chem. 1996;271(50):32253–32259. doi:10.1074/jbc.271.50.32253

32. Forsythe JA, Jiang BH, Iyer NV, et al. Activation of vascular endothelial growth factor gene transcription by hypoxia-inducible factor 1. Mol Cell Biol. 1996;16(9):4604–4613. doi:10.1128/MCB.16.9.4604

33. Gleadle JM, Ratcliffe PJ. Induction of hypoxia-inducible factor-1, erythropoietin, vascular endothelial growth factor, and glucose transporter-1 by hypoxia: evidence against a regulatory role for Src kinase. Blood. 1997;89(2):503–509.

34. Schuster SJ, Badiavas EV, Costa-Giomi P, Weinmann R, Erslev AJ, Caro J. Stimulation of Erythropoietin Gene Transcription During Hypoxia and Cobalt Exposure. Blood. 1989;73(1):13–16. doi:10.1182/blood.V73.1.13.13

35. Ebert BL, Bunn HF. Regulation of the Erythropoietin Gene. Blood. 1999;94(6):1864–1877. doi:10.1182/blood.V94.6.1864

36. Knyazev EN, Paul S. Levels of MIR-374 increase in BEWO B30 cells exposed to hypoxia [УРОВЕНЬ микроРНК MIR-374 ПОВЫШАЕТСЯ В КЛЕТКАХ BEWO B30 ПРИ ГИПОКСИИ]. Bull Russ State Med Univ. 2020;(2):11–17. doi:10.24075/BRSMU.2021.021

37. Strehl C, Fangradt M, Fearon U, Gaber T, Buttgereit F, Veale DJ. Hypoxia: how does the monocyte-macrophage system respond to changes in oxygen availability? J Leukoc Biol. 2014;95(2):233–241. doi:10.1189/jlb.1212627

38. Awad F, Assrawi E, Jumeau C, et al. Impact of human monocyte and macrophage polarization on NLR expression and NLRP3 inflammasome activation. PLOS ONE. 2017;12(4):e0175336. doi:10.1371/journal.pone.0175336

39. Sampath P, Moideen K, Ranganathan UD, Bethunaickan R. Monocyte Subsets: Phenotypes and Function in Tuberculosis Infection. Front Immunol. 2018;9:1726. doi:10.3389/fimmu.2018.01726

40. Vishnyakova P, Kuznetsova M, Poltavets A, et al. Distinct gene expression patterns for CD14++ and CD16++ monocytes in preeclampsia. Sci Rep. 2022;12(1):15469. doi:10.1038/s41598-022-19847-5

41. Tang MX, Zhang YH, Hu L, Kwak-Kim J, Liao AH. CD14++CD16+HLA-DR+ Monocytes in Peripheral Blood are Quantitatively Correlated with the Severity of Pre-eclampsia. Am J Reprod Immunol. 2015;74(2):116–122. doi:10.1111/aji.12389

42. Chen X, Wang Y, Qi Y, et al. Expansion of inflammatory monocytes in periphery and infiltrated into thyroid tissue in Graves’ disease. Sci Rep. 2021;11(1):13443. doi:10.1038/s41598-021-92737-4

43. Kokhanyuk B, Bodó K, Sétáló Jr G, Németh P, Engelmann P. Bacterial Engulfment Mechanism Is Strongly Conserved in Evolution Between Earthworm and Human Immune Cells. Front Immunol. 2021;12:733541. doi:10.3389/fimmu.2021.733541

44. Xiao J, Garcia-Lloret M, Winkler-Lowen B, Miller R, Simpson K, Guilbert LJ. ICAM-1-mediated adhesion of peripheral blood monocytes to the maternal surface of placental syncytiotrophoblasts: implications for placental villitis. Am J Pathol. 1997;150(5):1845–1860.

45. Kruger M, Coorevits L, De Wit TP, Casteels-Van Daele M, Van De Winkel JG, Ceuppens JL. Granulocyte-macrophage colony-stimulating factor antagonizes the transforming growth factor-beta-induced expression of Fc gamma RIII (CD16) on human monocytes. Immunology. 1996;87(1):162–167.

46. Korkes HA, De Oliveira L, Sass N, Salahuddin S, Karumanchi SA, Rajakumar A. Relationship between hypoxia and downstream pathogenic pathways in preeclampsia. Hypertens Pregnancy. 2017;36(2):145–150. doi:10.1080/10641955.2016.1259627

47. Zamudio S, Wu Y, Ietta F, et al. Human placental hypoxia-inducible factor-1alpha expression correlates with clinical outcomes in chronic hypoxia in vivo. Am J Pathol. 2007;170(6):2171–2179. doi:10.2353/ajpath.2007.061185

48. Keyes LE, Armaza FJ, Niermeyer S, Vargas E, Young DA, Moore LG. Intrauterine Growth Restriction, Preeclampsia, and Intrauterine Mortality at High Altitude in Bolivia. Pediatr Res. 2003;54(1):20–25. doi:10.1203/01.PDR.0000069846.64389.DC

49. Moore LG, Hershey DW, Jahnigen D, Bowes W. The incidence of pregnancy-induced hypertension is increased among Colorado residents at high altitude. Am J Obstet Gynecol. 1982;144(4):423–429. doi:10.1016/0002-9378(82)90248-4

50. Jones CJP, Fox H. An ultrastructural and ultrahistochemical study of the human placenta in maternal pre-eclampsia. Placenta. 1980;1(1):61–76. doi:10.1016/S0143-4004(80)80016-6

51. Ola MS, Nawaz Mohd, Ahsan H. Role of Bcl-2 family proteins and caspases in the regulation of apoptosis. Mol Cell Biochem. 2011;351(1):41–58. doi:10.1007/s11010-010-0709-x

52. Ratts VS, Tao XJ, Webster CB, et al. Expression of BCL-2, BAX and BAK in the Trophoblast Layer of the Term Human Placenta: a Unique Model of Apoptosis within a Syncytium. Placenta. 2000;21(4):361–366. doi:10.1053/plac.1999.0486

53. Nakashima A, Aoki A, Kusabiraki T, Cheng SB, Sharma S, Saito S. Autophagy regulation in preeclampsia: Pros and cons. J Reprod Immunol. 2017;123:17–23. doi:10.1016/j.jri.2017.08.006

54. Oh SY, Choi SJ, Kim KH, Cho EY, Kim JH, Roh CR. Autophagy-Related Proteins, LC3 and Beclin-1, in Placentas From Pregnancies Complicated by Preeclampsia. Reprod Sci. 2008;15(9):912–920. doi:10.1177/1933719108319159

55. Deleyto-Seldas N, Efeyan A. The mTOR–Autophagy Axis and the Control of Metabolism. Front Cell Dev Biol. 2021;9. Accessed June 13, 2023. https://www.frontiersin.org/articles/10.3389/fcell.2021.655731

56. Tsai K, Tullis B, Jensen T, Graff T, Reynolds P, Arroyo J. Differential expression of mTOR related molecules in the placenta from gestational diabetes mellitus (GDM), intrauterine growth restriction (IUGR) and preeclampsia patients. Reprod Biol. 2021;21(2):100503. doi:10.1016/j.repbio.2021.100503

57. Aiko Y, Askew DJ, Aramaki S, et al. Differential levels of amino acid transporters System L and ASCT2, and the mTOR protein in placenta of preeclampsia and IUGR. BMC Pregnancy Childbirth. 2014;14(1):181. doi:10.1186/1471-2393-14-181

58. Castellucci M, Kaufmann P. Basic Structure of the Villous Trees. In: Benirschke K, Kaufmann P, Baergen R, eds. Pathology of the Human Placenta. Springer; 2006:50–120. doi:10.1007/0-387-26742-5_6

59. de Luca Brunori I, Battini L, Brunori E, et al. Placental barrier breakage in preeclampsia: ultrastructural evidence. Eur J Obstet Gynecol Reprod Biol. 2005;118(2):182–189. doi:10.1016/j.ejogrb.2004.04.024

60. Anderson UD, Olsson MG, Rutardóttir S, et al. Fetal hemoglobin and α1-microglobulin as first- and early second-trimester predictive biomarkers for preeclampsia. Am J Obstet Gynecol. 2011;204(6):520.e1-520.e5. doi:10.1016/j.ajog.2011.01.058

61. Wong KL, Tai JJY, Wong WC, et al. Gene expression profiling reveals the defining features of the classical, intermediate, and nonclassical human monocyte subsets. Blood. 2011;118(5):e16–e31. doi:10.1182/blood-2010-12-326355

62. Prabhu VM, Singh AK, Padwal V, Nagar V, Patil P, Patel V. Monocyte Based Correlates of Immune Activation and Viremia in HIV-Infected Long-Term Non-Progressors. Front Immunol. 2019;10. Accessed June 14, 2023. https://www.frontiersin.org/articles/10.3389/fimmu.2019.02849

63. Lehman N, Kowalska W, Zarobkiewicz M, et al. Pro- vs. Anti-Inflammatory Features of Monocyte Subsets in Glioma Patients. Int J Mol Sci. 2023;24(3):1879. doi:10.3390/ijms24031879

64. Chuluundorj D, Harding SA, Abernethy D, La Flamme AC. Expansion and preferential activation of the CD14+CD16+ monocyte subset during multiple sclerosis. Immunol Cell Biol. 2014;92(6):509–517. doi:10.1038/icb.2014.15

65. Ong SM, Hadadi E, Dang TM, et al. The pro-inflammatory phenotype of the human non-classical monocyte subset is attributed to senescence. Cell Death Dis. 2018;9(3):1–12. doi:10.1038/s41419-018-0327-1

66. Thomas G, Tacke R, Hedrick CC, Hanna RN. Nonclassical patrolling monocyte function in the vasculature. Arterioscler Thromb Vasc Biol. 2015;35(6):1306–1316. doi:10.1161/ATVBAHA.114.304650

67. Flood-Nichols SK, Kazanjian AA, Tinnemore D, et al. Aberrant Glycosylation of Plasma Proteins in Severe Preeclampsia Promotes Monocyte Adhesion. Reprod Sci. 2014;21(2):204–214. doi:10.1177/1933719113492210

