## Supplemental Table and Figures for "Bi-directional communication between monocytes and trophoblasts under hypoxia and hypoxia-reperfusion conditions"

**Supplemental Table 1: Placenta IDs and Limited Patient Demographics**

| **Placenta ID:** | **Gestational Age:** | **Complications:** |
| --- | --- | --- |
| 1 | 31w4d | PE with severe features |
| 3 | 30w3d | Intrauterine growth restriction |
| 4 | 32w3d | Placental Abruption |
| 8 | 32w3d | PE with severe features |
| 9 | 36w5d | Placenta previa |
| 12 | 37w0d | None |
| 18 | 37w0d | Gestational hypertension |
| 21 | 34w5d | Vasa previa |
| 22 | 34w1d | Twins, IUGR |
| 25 | 37w0d | PE without severe features |
| 29 | 33w6d | PE with severe features |
| 31 | 39w0d | None |
| 34 | 31w2d | PE with severe features |
| 37 | 35w1d | Vasa previa |
| 38 | 34w1d | PE with severe features |
| 39 | 35w0d | Vasa previa |

**Supplemental Table 2: qPCR Primers**

| **Gene Name:** | **Abbreviation:** | **NCBI Gene ID:** | **Sequence:** |
| --- | --- | --- | --- |
| Actin-β | ACTB | 60 | F: CCCTGGACTTCGAGCAAGAG  R: ACTCCATGCCCAGGAAGGAA |
| Hypoxia-Inducible Factor 1-α | HIF-1α | 3091 | F: CACCACAGGACAGTACAGGAT  R: CGTGCTGAATAATACCACTCACA |
| Erythropoietin | EPO | 2056 | F: GGAGGCCGAGAATATCACGAC  R: CCCTGCCAGACTTCTACGG |
| Glucose Transporter-1 | GLUT1 | 6513 | F: GGCCAAGAGTGTGCTAAAGAA  R: ACAGCGTTGATGCCAGACAG |
| Vascular Endothelial Growth Factor | VEGF | 7422 | F: AGGGCAGAATCATCACGAAGT  R: AGGGTCTCGATTGGATGGCA |
| Binding of Immunoglobulin Protein | BiP | 3309 | F: CATCACGCCGTCCTATGTCG  R: CGTCAAAGACCGTGTTCTCG |
| CCAAT/Enhancer-Binding Protein Homologous Protein | Chop | 1649 | F: GAACGGCTCAAGCAGGAAATC  R: TTCACCATTCGGTCAATCAGAG |
| BcL2-Associated X-Protein | Bax | 581 | F: GACCAGCATGACAGATTTCTACCA  R: AACTGAGACTAAGGCAGAAGATG |
| BcL2 | - | 596 | F: TCCCTCGCTGCACAAATACTC  R: TTCTGCCCCTGCCAAATCT |
| Microtubule-Associated Protein 1A/1B-Light Chain 3 | LC3 | 84577 | F: AAGGCGCTTACAGCTCAATG  R: CTGGGAGGCATAGACCATGT |
| Mammalian Target of Rapamycin | mTOR | 2475 | F: ATGCTTGGAACCGGACCTG  R: TCTTGACTCATCTCTCGGAGTT |
| C-X3-C Motif Chemokine Ligand 1 | CX3CL1 | 6376 | F: ACCACGGTGTGACGAAATG  R: TGTTGATAGTGGATGAGCAAAGC |
| Intercellular Adhesion Molecule 1 | ICAM-1 | 3383 | F: ATGCCCAGACATCTGTGTCC  R: GGGGTCTCTATGCCCAACAA |
| Sialic-Acid Binding Ig-like lectin 3 | CD33 | 945 | F: GGCCACTCCAAAAACCTGAC  R: GACAACCAGGAGAAGATCGGG |
| Cluster of Differentiation 40 | CD40 | 958 | F: ACTGAAACGGAATGCCTTCCT  R: CCTCACTCGTACAGTGCCA |
| Cluster of Differentiation 64 | CD64 | 2209 | F: TGGGTCAGCGTGTTCCAAG  R: CACCTGTATTCACCACTGTCATT |
| Cluster of Differentiation 86 | CD86 | 942 | F: CTGCTCATCTATACACGGTTACC  R: GGAAACGTCGTACAGTTCTGTG |
| Cluster of Differentiation 163 | CD163 | 9332 | F: CCAGAAGGAACTTGTAGCCACAG  R: CAGGCACCAAGCGTTTTGAGCT |
| Mannose Receptor C-Type 1 | CD206 | 4360 | F: TCCGGGTGCTGTTCTCCTA  R: CCAGTCTGTTTTTGATGGCACT |
| C-X-C Motif Chemokine Receptor 1 | CXCR1 | 3577 | F: CTGACCCAGAAGCGTCACTTG  R: CCAGGACCTCATAGCAAACTG |
| C-X-C Motif Chemokine Receptor 2 | CXCR2 | 3579 | F: CCTGTCTTACTTTTCCGAAGGAC  R: TTGCTGTATTGTTGCCCATGT |
| C-C Motif Chemokine Receptor 5 | CCR5 | 1234 | F: GTTGGACCAAGCTATGCAGGT  R: GCAGAAGCGTTTGGCAATGT |
| Major Histocompatibility Complex, Class II, DR alpha | HLA-DRα | 3122 | F: AGTCCCTGTGCTAGGATTTTTCA  R: ACATAAACTCGCCTGATTGGTC |
| Major Histocompatibility Complex, Class II, DR beta 1 | HLA-DRβ1 | 3123 | F: CGGGGTTGGTGAGAGCTTC  R: AACCACCTGACTTCAATGCTG |
| Major Histocompatibility Complex, Class II, DR beta 3 | HLA-DRβ3 | 3125 | F: CGGGGTTGGTGAGAGCTTC  R: AACCACCTGACTTCAATGCTG |
| C-C Motif Chemokine Receptor 2 | CCR2 | 729230 | F: CCACATCTCGTTCTCGGTTTATC  R: CAGGGAGCACCGTAATCATAATC |
| CX3C Motif Chemokine Receptor 1 | CX3CR1 | 1524 | F: ACTTTGAGTACGATGATTTGGCT  R: GGTAAATGTCGGTGACACTCTT |
| Leukocyte Function-Associated Antigen 1 | LFA-1 | 3683 | F: TGCTTATCATCATCACGGATGG  R: CTCTCCTTGGTCTGAAAATGCT |
| Interleukin-8 | IL-8 | 3576 | F: TTTTGCCAAGGAGTGCTAAAGA  R: AACCCTCTGCACCCAGTTTTC |
| Interleukin-10 | IL-10 | 3386 | F: TCTCCGAGATGCCTTCAGCAGA  R: TCAGACAAGGCTTGGCAACCCA |
| Interleukin-12 | IL-12 | 3592, 3593 | F: TGCCTTCACCACTCCCAAAACC  R: CAATCTCTTCAGAAGTGCAAGGG |
| Tumor Necrosis Factor | TNF-α | 7124 | F: CCTCTCTCTAATCAGCCCTCTG  R: GAGGACCTGGGAGTAGATGAG |
| Transforming Growth Factor Beta 1 | TGF-β1 | 7040 | F: GGCCAGATCCTGTCCAAGC  R: GTGGGTTTCCACCATTAGCAC |
| C-C Motif Chemokine Ligand 3 | MIP-1α | 6348 | F: AGTTCTCTGCATCACTTGCTG  R: CGGCTTCGCTTGGTTAGGAA |
| Nitric Oxide Synthase 2 | NOS2 | 4843 | F: GCTCTACACCTCCAATGTGACC  R: CTGCCGAGATTTGAGCCTCATG |

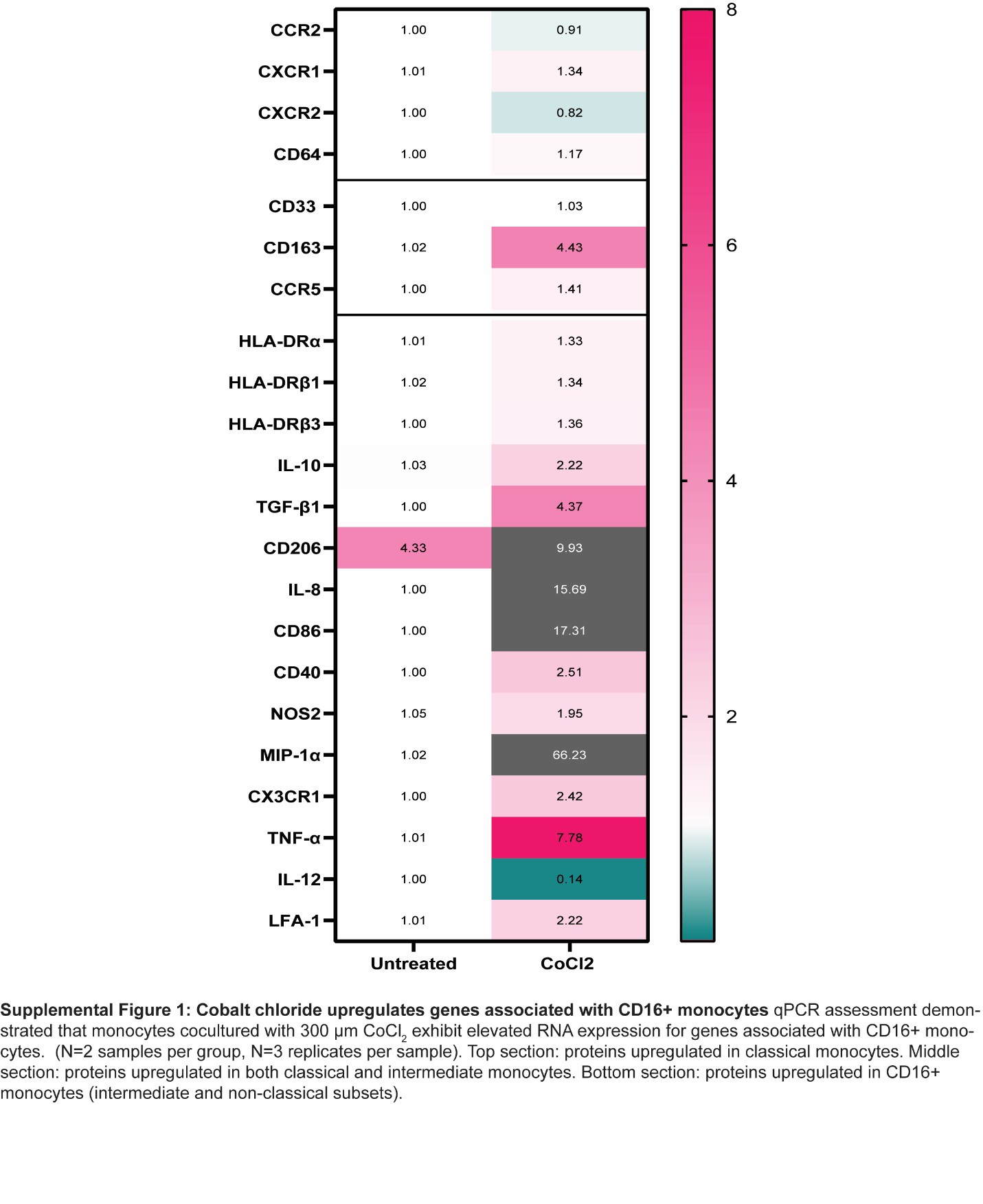
